# Uneven growth of SARS-CoV-2 clones evidenced by more than 500,000 whole-genome sequences

**DOI:** 10.1101/2021.04.06.437914

**Authors:** Hong-Li Zeng, Yue Liu, Kaisa Thorell, Rickard Nordén, Erik Aurell

## Abstract

We have computed the frequencies of the alleles of the “UK variant” (B.1.1.7) and “South Africa variant” (B.1.351) of SARS-CoV-2 from the large GISAID repository. We find that the frequencies of the mutations in UK variant overall rose towards the end of 2020, as widely reported in the literature and in the general press. However, we also find that these frequencies vary in different patterns rather than in concert. For South Africa variant we find a more complex scenario with frequencies of some mutations rising and some remaining close to zero. Our results point to that what is generally reported as one variant is in fact a collection of variants with different genetic characteristics.

## Introduction

COVID-19 has so far led to the confirmed deaths of more than 2,700,000 people (*1*) and has caused the largest disruption in the world economy and human life for several generations. While several efficient vaccines have been developed and some countries have already progressed far towards herd immunity, most of the world is still in the midst of the pandemic. As its elimination in many countries will likely only happen on the time scales of years and not months, a better understanding of the biology of SARS-CoV-2 will remain of high importance.

The GISAID repository (*2*) contains a rapidly increasing collection of SARS-CoV-2 whole-genome sequences, and has been used to identify mutational hotspots and potential drug targets (*3*) as well as to infer epistatic fitness parameters (*4*, *5*). In recent months Nature has performed an experiment in the growth of Variants of Concern (VOC) B.1.1.7 and B.1.351. These variants of the virus are commonly referred to as “UK variant” and “South Africa variant”, as they were first identified in south-east England (*6*), and South Africa (*7*, *8*) respectively. Although most assays for these variants are based on variability at a few positions, the full original definitions contain more loci, both in Spike and outside Spike. For UK variant the original definition contains mutations at 23 positions well spread out along the SARS-CoV-2 genome. The frequencies of the mutations at the different positions hence give information on whether these variants in fact grow as large clones, or if they have mutated or recombined into several clones, or if they were several clones from the beginning. We find that the second scenario holds for UK variant while a combination of the second and third scenario holds for South Africa variant.

## Results

The GISAID repository (*2*) holds a large collection of SARS-CoV-2 whole-genome sequences. In the following we have used genomes qualified as “high quality” and annotated with sampling date up to the end of February, 2021. We note that submission date to GISAID is later than sampling date, typically by two weeks or more. The data used hence represents a large part of all the whole-genome sequences available on GISAID up to mid-March 2021. The total number of SARS-CoV-2 genomes used in this study is 562,477. The data has been stratified by sampling time, as shown in figure captions.

The first report from Public Health England (Technical briefing 1, December 21, 2020) defining B.1.1.7 as a Variant of Concern lists 17 non-synonymous mutations. (including deletions) and six synonymous mutations (*6*), see Materials & Methods Table 1. Of these 23 mutations, 21 have a similar time course in time-sorted GISAID data, *C*16176*T* has the precise opposite time course, and *T*26801*C* an unrelated time course, see Fig. 1. In the following we have assumed that *C*16176*T* is a mis-labelling, and that this mutation in fact is *T*16176*C*. We have further assumed that *T*26801*C*, a synonymous mutation in the *M* gene, pertains mostly to another clone or to another reference sequence. In the following we have not retained data from this locus.

**Table 1:**
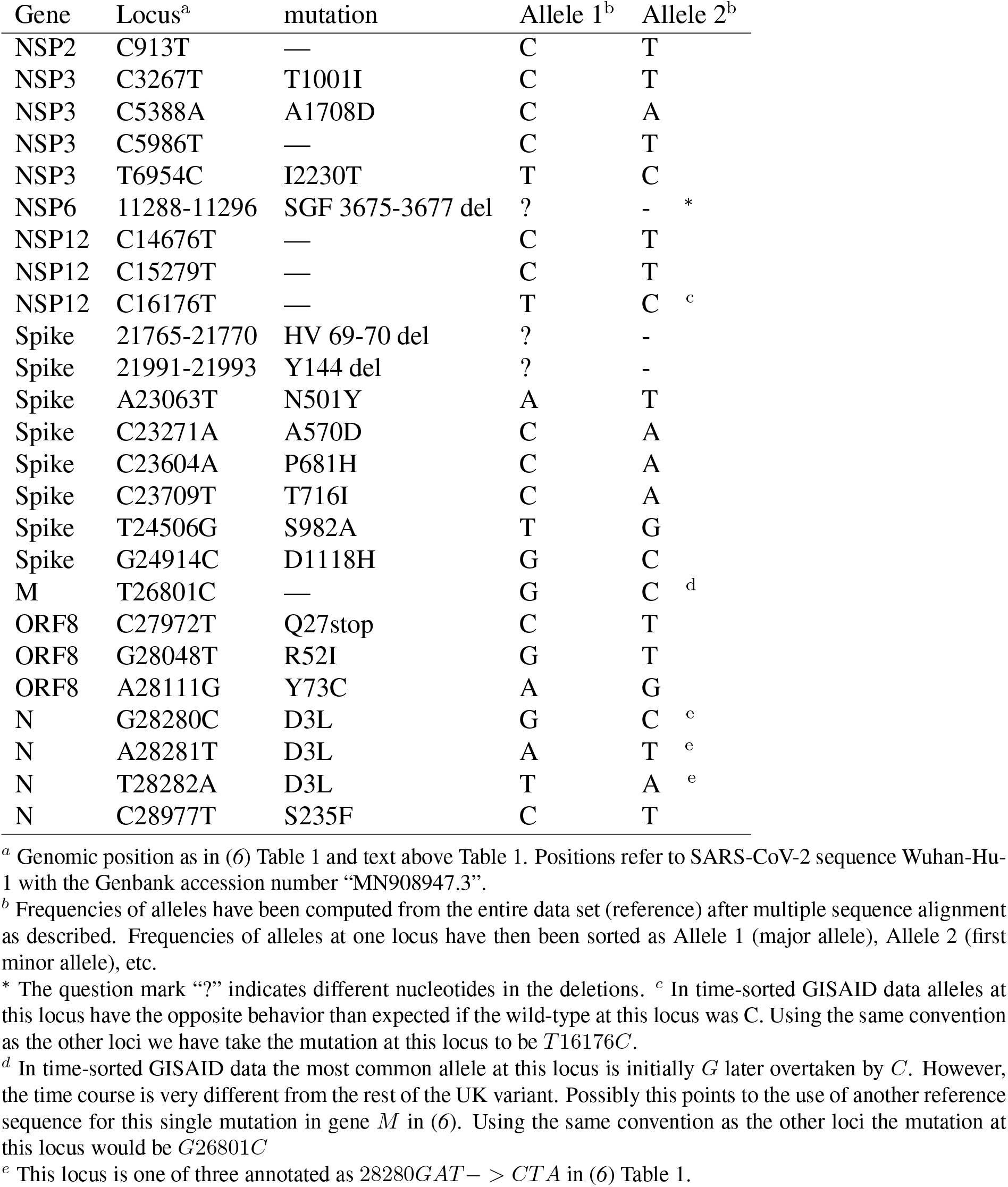
Defining mutations for variant B.1.1.7

| Gene | Locus<sup>a</sup> | mutation | Allele 1<sup>b</sup> | Allele 2<sup>b</sup> |
| --- | --- | --- | --- | --- |
| NSP2 | C913T | — | C | T |
| NSP3 | C3267T | T1001I | C | T |
| NSP3 | C5388A | A1708D | C | A |
| NSP3 | C5986T | — | C | T |
| NSP3 | T6954C | I2230T | T | C |
| NSP6 | 11288-11296 | SGF 3675-3677 del | ? | - * |
| NSP12 | C14676T | — | C | T |
| NSP12 | C15279T | — | C | T |
| NSP12 | C16176T | — | T | C <sup>c</sup> |
| Spike | 21765-21770 | HV 69-70 del | ? | - |
| Spike | 21991-21993 | Y144 del | ? | - |
| Spike | A23063T | N501Y | A | T |
| Spike | C23271A | A570D | C | A |
| Spike | C23604A | P681H | C | A |
| Spike | C23709T | T716I | C | A |
| Spike | T24506G | S982A | T | G |
| Spike | G24914C | D1118H | G | C |
| M | T26801C | — | G | C <sup>d</sup> |
| ORF8 | C27972T | Q27stop | C | T |
| ORF8 | G28048T | R52I | G | T |
| ORF8 | A28111G | Y73C | A | G |
| N | G28280C | D3L | G | C <sup>e</sup> |
| N | A28281T | D3L | A | T <sup>e</sup> |
| N | T28282A | D3L | T | A <sup>e</sup> |
| N | C28977T | S235F | C | T |

<sup>a</sup> Genomic position as in (6) Table 1 and text above Table 1. Positions refer to SARS-CoV-2 sequence Wuhan-Hu-1 with the Genbank accession number “MN908947.3”.
<sup>b</sup> Frequencies of alleles have been computed from the entire data set (reference) after multiple sequence alignment as described. Frequencies of alleles at one locus have then been sorted as Allele 1 (major allele), Allele 2 (first minor allele), etc.
\* The question mark “?” indicates different nucleotides in the deletions. <sup>c</sup> In time-sorted GISAID data alleles at this locus have the opposite behavior than expected if the wild-type at this locus was C. Using the same convention as the other loci we have take the mutation at this locus to be *T16176C*.
<sup>d</sup> In time-sorted GISAID data the most common allele at this locus is initially *G* later overtaken by *C*. However, the time course is very different from the rest of the UK variant. Possibly this points to the use of another reference sequence for this single mutation in gene *M* in (6). Using the same convention as the other loci the mutation at this locus would be *G26801C*
<sup>e</sup> This locus is one of three annotated as *28280GAT* → *CTA* in (6) Table 1.

**Figure 1:**
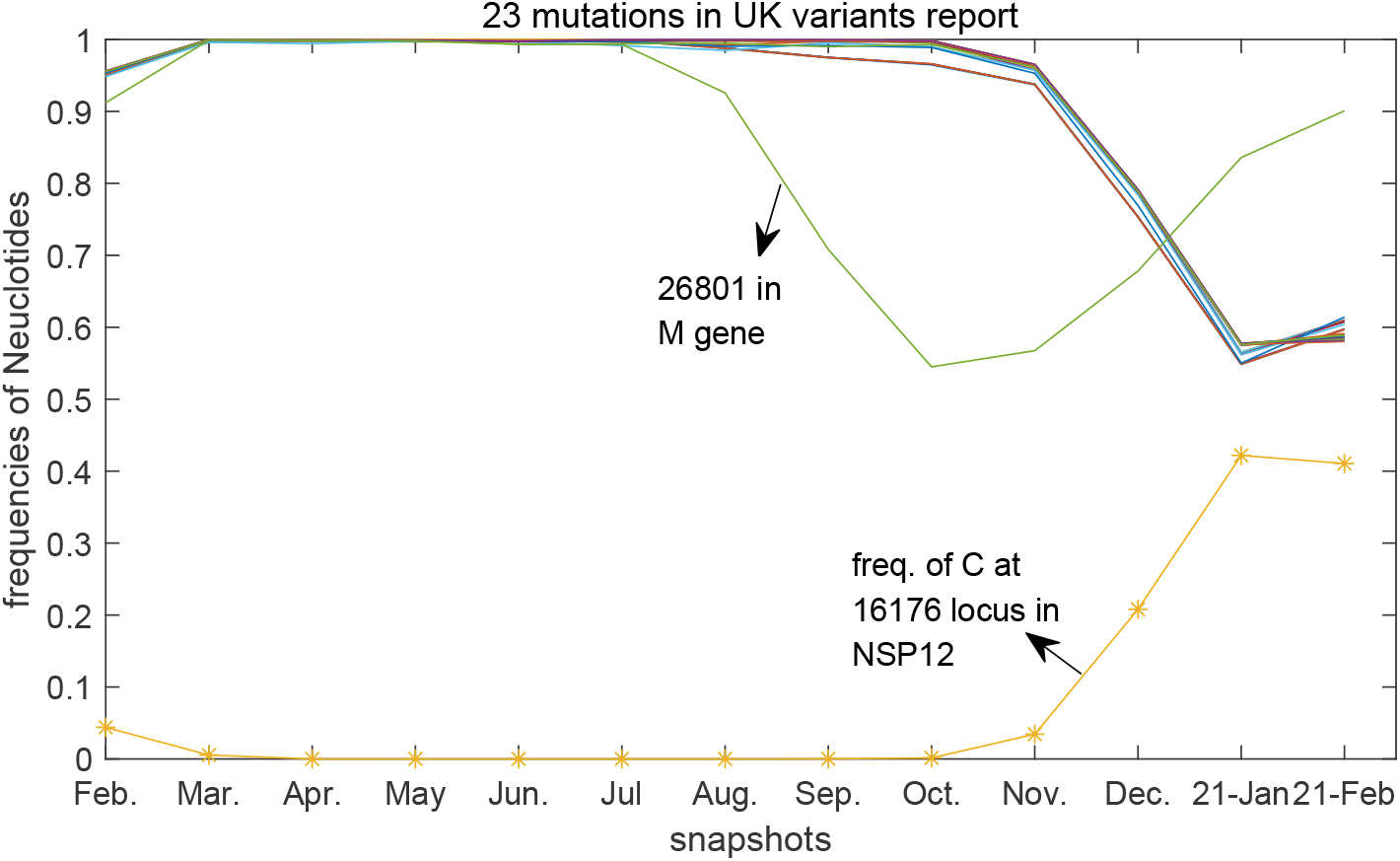
Frequency of major allele at the defining loci for B.1.1.7 over time as determined from GISAID. 21 out of 23 mutations listed for the UK variant report have similar temporal pattern, except the 26801 locus in M gene and the mutation of C16176T in NSP12.

The definition of B.1.351 given by Public Health England (Technical briefing 6, February 13, 2020) lists 17 non-synonymous mutations (including deletions) out of which nine in Spike (*8*), see Materials & Methods Table 2. Of these 17 mutations, three appeared much before this variant was defined and have an unrelated time course, see Fig. 2. In the following we have assumed that these three mutations, *C*1059*T* (*T* 265*I* in *NSP* 2), *C*21614*T* (*L*18*F* in Spike) and *G*25563*T* (*Q*57*H* in *ORF* 3*a*) mostly pertain to other clones and/or to another reference sequence. We have not retained data from these loci. The frequencies of the other loci include two that are also present in B.1.1.7 and follow that course, and the rest which remain at an order-of-magnitude lower level in the GISAID data used here.

**Table 2:** Defining mutations for variant B.1.351

| Gene | Locus<sup>a</sup> | mutation | Allele 1<sup>b</sup> | Allele 2<sup>b</sup> |
| --- | --- | --- | --- | --- |
| NSP2 | C1059T | T265I | C | T |
| NSP3 | G5230T | K1655N | G | T<sup>d</sup> |
| NSP5 | A10323G | K3353R | A | G |
| NSP6 | 11288_96 del | 3675-3677 del | — | —<sup>e</sup> |
| Spike | C21614T | L18F | C | T<sup>c</sup> |
| Spike | A21801C | D80A | A | C<sup>d</sup> |
| Spike | A22206G | D215G | A | G<sup>d</sup> |
| Spike | — | 242-244del | — | — |
| Spike | G22299T | R246I | G | T<sup>c</sup> |
| Spike | G22813T | K417N | G | T<sup>c,d</sup> |
| Spike | G23012A | SGF E484K | G | A<sup>d</sup> |
| Spike | A23063T | N501Y | A | T<sup>d,e</sup> |
| Spike | C23664T | A701V | C | T<sup>d</sup> |
| ORF3a | G25563T | Q57H | G | T |
| ORF3a | C25904T | S171L | C | T |
| E | C26456T | P71L | C | T<sup>d</sup> |
| N | C28887T | T205I | C | T<sup>d</sup> |

<sup>a</sup> Genomic position as in (8) Table 4a. Positions refer to SARS-CoV-2 sequence Wuhan-Hu-1 with the Genbank accession number “MN908947.3”.
<sup>b</sup> Frequencies of alleles have been computed from the entire data set (reference) after multiple sequence alignment as described. Frequencies of alleles at one locus have then been sorted as Allele 1 (major allele), Allele 2 (first minor allele), etc.
<sup>c</sup> Annotated in (8) caption to Table 4a as acquisitions in subset of isolates within the lineage.
<sup>d</sup> Annotated in (8) in Table 4b as “PROBABLE”; at least 4 lineage defining non-synonymous changes called as alternate base and all other positions either N or mixed base OR at least 5 of the 9 non-synonymous changes.
<sup>e</sup> This mutation is also present in the UK variant, compare Table 1.

**Figure 2:**
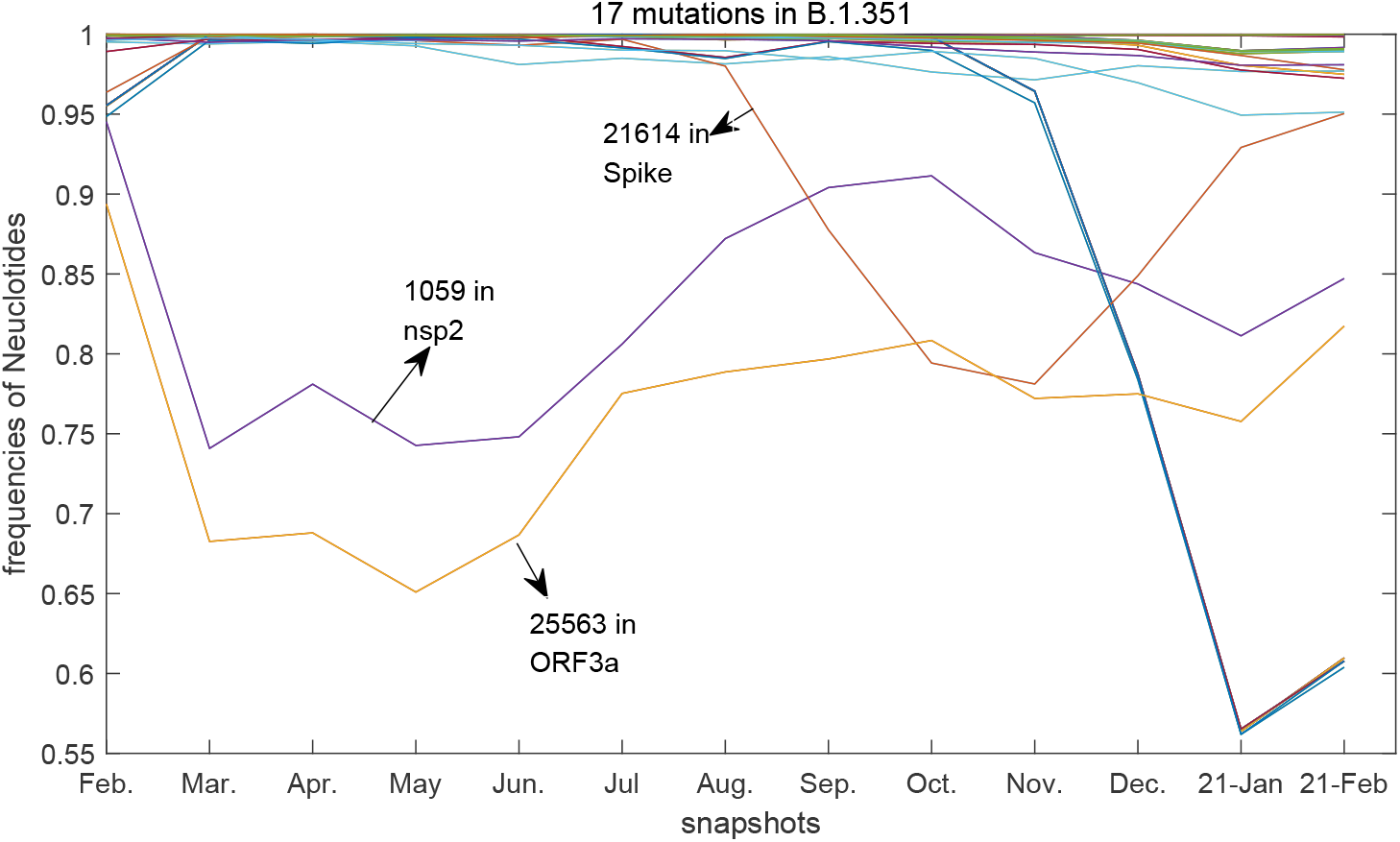
Frequency of major allele at the defining loci for B.1.351 over time as determined from GISAID. Three out of 17 mutations listed for the South Africa variant (marked in figure) display a different dynamics and have been excluded from the following analysis. Of the others, two mutations shared with B.1.1.7 increase to large frequencies: the 3675-3677 deletion (11288-11296) in NSP6 and the N501Y mutation (*A*23063*T*) in Spike. The remaining mutations reach about the 2% level and are discussed in text.

The frequencies of the 22 retained mutations for the UK variant increase in frequency after late summer / early autumn 2020, see Fig. 3. The lines in this figure connect frequencies of the second most common allele (first minor allele) within the same month of sampling time in the GISAID data. With one exception (16176, discussed above) this second most common allele agrees with the mutation at this locus as given in (*6*).

**Figure 3:**
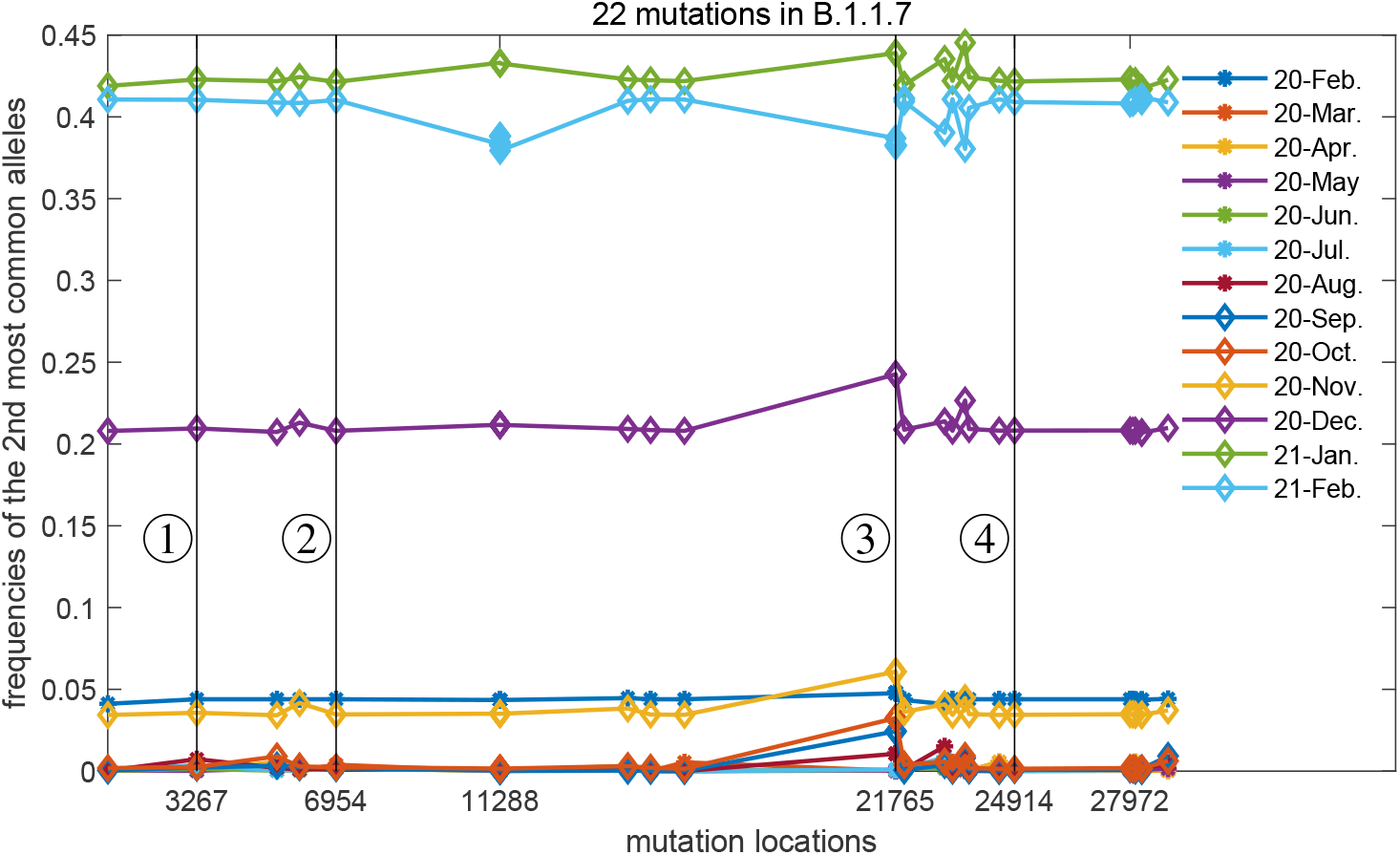
Frequency of second most common allele at the defining loci for B.1.1.7 over time as determined from GISAID. x-axis gives genomic position. The 26801 locus in *M* gene is not included compared with Fig. 1. The first and second vertical lines indicates the main non-synonymous mutations in NSP3 while the third and fourth ones mark the B.1.1.7 mutations in Spike. The average distance between the 22 retained mutations is about 1, 500bp, but some such as *C*23604*A* and *C*23709*T* (*P*681*H* and *T*716*I* in Spike) lie closer. The sixth mutation from the left (first to the right of line marked 2 is the SGF 3675-3677 deletion in NSP6 (11288-11296). The tenth mutation from the left (at line marked 3) is the HV 69-70 deletion in Spike (21765-21770).

**Figure 4:**
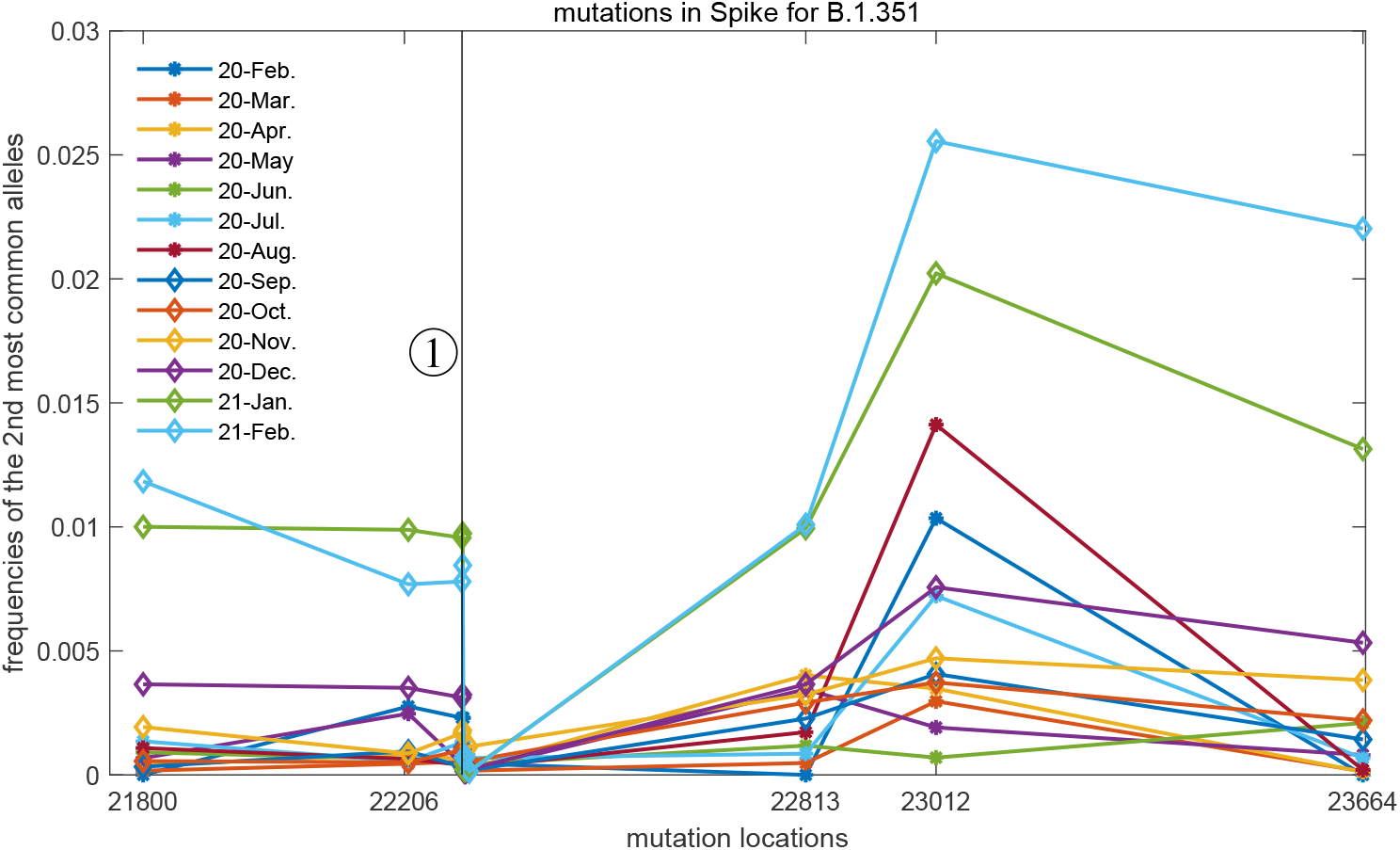
Frequency of second most common allele at the defining loci in Spike for B.1.351 over time as determined from GISAID. x-axis gives genomic position. The line marked 1 indicates two very closely spaced mutations in Spike S1, the 242-244 deletion (immediately to the left) and R246I (*G*22299*T*, immediately to the right). The mutation at 23012 is SGF E484K in Spike S2. The mutation N501Y (*A*23063*T*) is also defined for B.1.1.7 and is not shown in the above, see instead Fig. 3.

The growth of the first minor allele of the UK variant is uneven across the SARS-CoV-2 genome. In a first phase (early 2020-November 2020), the frequency of the HV 69-70 deletion in Spike (21765-21770) is noticeably higher than the other mutations defining B.1.1.7. This is consistent with this mutation initially being present also in clones unrelated to B.1.1.7. As time progresses, the relative difference between the frequency at this locus and the frequencies at the other loci decreases. In December 2020 the frequency of *C*23604*A* (*P*681*H* in Spike) also noticeably increases above the others. For the last two months (January and February 2021) one further observes that the frequencies of the deletion 11288 – 11296 in *NSP* 6 and the mutation *A*23063*T* (*N*501*Y* in Spike) to be noticeably different from the others.

For the South Africa variant we have chosen to focus on mutations in Spike, except N501Y in Spike (*A*23063*T*) which is shared with the UK variant, see Fig. 3.

The growth of these mutations in B.1.351 in Spike are as follows. From the beginning of Spike up and including the 242-244 deletion (three loci) there is a roughly even growth up to approximately the 1% level. This means that in January and February 2021 about 1% of the SARS-CoV-2 genomes sequenced world-wide carried these mutations. Immediately to the right there is a sharp drop in frequency so that very few sequences carried the R246I mutation. K417N (*G*22813*T*) and A701V (*C*23664*T*) follows approximately the pattern of the 242-244 deletion albeit A701V appears to grow faster in frequency towards the end of period. E484K (*G*23012*A*) on the other hand follows an erratic trajectory peaking at above 1% in August 2020, falling back to below 1% in November 2020, and then increasing again up to 2.0 – 2.5% in January-February 2021.

## Materials and Methods

### Data Collection

We analyzed the consensus sequences deposited in the GISAID database (*2*) (https://www.gisaid.org) with high quality and full lengths (number of bps ≈ 30,000) which can be obtained through the options of “complete” and “high coverage” on the GISAID interface. The “collection time” option was clicked for the convenience of data analysis in the following steps. The sequences were downloaded from GISAID website around the middle of March 2021 setting collection time until the end of February 2021. The total number of selected genomic sequences was 562, 477. The sequences are listed and available on the Github repository (*9*).

### Multiple-Sequence Alignment (MSA)

Multiple sequence alignments (MSAs) were constructed with the online alignment server MAFFT (*10*, *11*) with the reference sequence “Wuhan-Hu-1” with the Genbank accession number “MN908947.3”. The length of sequences are kept the same as the reference during sequence alignment.

An MSA is a big matrix 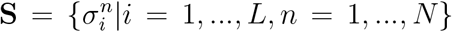, composed of *N* genomic sequences which are aligned over *L* positions. Each entry 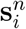 of matrix **S** is either one of the 4 nucleotides (A,C,G,T), or “not known nucleotide” (N), or some minorities, or the alignment gap ‘-’ introduced to treat nucleotide deletions or insertions,. The minorities ‘KFY…’ are transformed into ‘N’ in the process of constructing the MSA. Then there are hence six remaining states, i.e., ‘-NACGT’.

To reduce the burden of the desktop computer, the whole MSA are saved in forms of each sub-structures (NSP1 to NSP16, Spike, ORF3a and other genes) for the SARS-CoV-2 sequences. Furthermore, the data for the whole 2020 year is stored in one data file while the data for January and February 2021 are in two separate data files as they contain 114,858 and 99,518 sequences respectively.

### Data Storage

The SARS-CoV-2 dataset downloaded from GISAID website are stored in a desktop computer with 64G RAM named “hlz” at Nanjing University of Posts and Telecommunications (NJUPT).

### Frequency computations and visualizations

The allele frequencies and the visualizations were both done using MATLAB R2020a on “hlz”.

### Data Analysis

The work mainly focused on the allele frequencies analysis for the mutations or deletions listed for B.1.1.7 (*6*) also known as “UK variant” and B.1.351 (*8*) also known as “South Africa variant”. For a certain time period Δ*t*, the frequencies of a certain nucleotide *x* at *i* locus are computed by eq. 1.

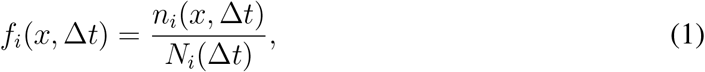

With *x* ∈ {–*, N, A, C, G, T*} and Δ*t* the time length of the analyzed snapshots. *n_i_*(*x,* Δ*t*) denotes the number of allele *x* at locus *i* during the period of Δ*t* while the denominator is the total number of the nucleotides on this locus during the same period Δ*t*.

To take into account the effect of evolution time for SARS-CoV-2 virus, the allele frequencies are computed on the time scale of each month from the initial outbreak of the COVID-19 pandemic. The outliers pointed by arrows in Fig. 1 and Fig. 2 are identified manually.

### Annotated nucleotide mutations

Differently from the allele frequencies based on the data snapshots (Months), the annotated nucleotide mutations are obtained from the whole dataset. With the sorted allele frequencies computed from the whole dataset, the most prevalent nucleotide and the second most one are selected as the first and second allele in the mutations shown in Tab. 1 and Tab. 2.

### Definition of B.1.1.7 (“UK variant”)

In this work we have used the definition of SARS-CoV-2 Variant of Concern 202012/01 (B.1.1.7) as originally given in “Technical briefing 1” (*6*) Table 1 and text above Table 1 (publication date December 21, 2020). This information with annotations is given as Table 1 below.

In a later report from the same group (Technical briefing 6, publication date February 13, 2021 (*8*)) another definition of B.1.1.7 was given in Table 2a. That definition differs from the one used here in that mutation *C*28977*T* in the *N* gene and the six synonymous mutations have not been retained.

### Definition of B.1.351 (“South Africa variant”)

In this work we have used the definition of SARS-CoV-2 Variant of Concern 202012/02 (B.1.351) as given in “Technical briefing 6” (*8*) Table 4a. This information with annotations is given as Table 2 below.

## Discussion

The conclusion of this work is that it cannot be the case that UK variant and South Africa variant, as originally defined, grow as two large clones. Disregarding the two mutations in B.1.1.7 and three mutations in B.1.351 with clearly deviating time series behaviour, the dynamics at the other loci is sufficiently different to rule out one-clone scenarios. No sophisticated statistical analysis is required to reach this conclusion. In this work we have used well over half a million whole-genome SARS-CoV-2 sequences from GISAID, and for the last two months the plots of the monthly frequency data in above are based on the order of 100, 000 sequences.

The instability of clones is supported by recent observations points towards the emergence of multiple lineages of SARS-CoV-2 within the same individual (*12*, *13*, *14*, *15*, *16*). In all cases the patients had prolonged viremia and received convalescent plasma treatment and/or monoclonal antibody therapy. Treatment with convalescent plasma or monoclonal antibodies applies selection pressure on a viral population within the host that may drive the emergence of antibody resistant clones. Also, the large number of viral genomes present simultaneously in a single patient enable opportunities for within host recombination. The phenotypic effects of all described mutations in the spike protein of SARS-CoV-2 are just beginning to be unraveled. For example, the N501Y substitution increases the affinity for ACE2 binding (*17*). Also, compensatory mutations have been described as in the case for the E484K substitution in combination with del69-70, where a reduction in antibody sensitivity is compensated with increased infectivity.

Coronaviruses, the larger family to which SARS-CoV-2 belongs, in general exhibit a large amount of recombination (*18*, *19*). There are reports that this is so also for SARS-CoV-2 (*20*, *21*, *22*). Large-scale recombination would be important in the COVID19 pandemic for several reasons. First it increases the resiliance of the viral population against hostile agents. Beneficial (to the virus) changes can spread faster and more reliably throughout the population. Second it leads to form of evolution optimizing fitness and less impacted by traits inherited by chance. While a clone replicating asexually will likely have points of weakness, in a recombining population such errors are shared around and eliminated. Third, substantial amount of recombination is a confounder for phylogentic reconstruction. Crudely put, phylogenetic trees reconstructed from population-wide sequence data may not reflect the actual evolution in such populations, an issue which has been discussed in bacterial phylogenetics since some time (*23*, *24*, *25*). Lastly, a population under strong recombination is expected to be in Kimura’s Quasi-Linkage Equilibrium (*26*, *27*, *28*) which allows efficient and accurate inference of evolutionary parameters from sequence information (*4*, *5*). On a positive note this opens up the perspective of systematic search for new drugs and combinatorial drug treatments by leveraging large-scale whole-genome sequencing data.

## Acknowledgments

We thank Richard Neher for comments on a first version of the MS, and for pointing out that in the annotation used by nextstrain, *C*26801*G* is counted in clade 20E (EU1). The work of HLZ was sponsored by National Natural Science Foundation of China (11705097). The work of EA was supported by the Swedish Research Council grant 2020-04980.

